# DYNAMICS OF NEUROMUSCULAR TRANSMISSION REPRODUCED BY CALCIUM-DEPENDENT SERIAL TRANSITIONS IN THE VESICLE FUSION COMPLEX

**DOI:** 10.1101/2021.09.15.460525

**Authors:** Alejandro Martínez-Valencia, Guillermo Ramírez-Santiago, Francisco F. De-Miguel

## Abstract

Neuromuscular transmission, from spontaneous release to facilitation and depression was accurately reproduced by a mechanistic kinetic model of sequential maturation transitions in the molecular fusion complex. The model incorporates three predictions. First, sequential calcium-dependent forward transitions take vesicles from docked to pre-primed to primed states, followed by fusion. Second, pre-priming and priming are reversible. Third, fusion and recycling are unidirectional. The model was fed with experimental data from previous studies while the backward (*β*) and recycling (*ρ*) rate constant values were fitted. Classical experiments were successfully reproduced when every forward (*α*) rate constant had the same value, and both backward rate constants were 50-100 times larger. Such disproportion originated an abruptly decreasing gradient of resting vesicles from docked to primed states. Simulations also predict that: i. Spontaneous release reflects primed to fusion spontaneous transitions. ii. Calcium elevations synchronize the series of forward transitions that lead to fusion. iii Facilitation reflects a transient increase of priming following calcium-dependent transitions. iv. Backward transitions and recycling restore the resting state. v. Depression reflects backward transitions and slow recycling after intense release. Such finely-tuned kinetics offers a mechanism for collective non-linear transitional adaptations of a homogeneous vesicle pool to an ever-changing pattern of electrical activity.

## Introduction

In the present study we searched for a unifying molecular mechanism by which neuromuscular transmission adapts dynamically to the ongoing pattern of electrical activity. Four aspects of transmission were analyzed in detail. i. Spontaneous release at rest (Fatt and Katz, 1952); ii. Calcium-dependent evoked release on an impulse (Katz and Miledi, 1979), iii. Facilitation, namely a non-linear increase of release upon rapid subsequent stimulation (Feng, 1940; Eccles, 1941; Liley and North, 1953; del Castillo and Katz, 1954b: Katz and Miledi, 1968), and iv. Depression, namely a reduction of release on stimulation at extended intervals under high release probability (Eccles, Katz, Kuffler, 1941; Lundberg and Quilish, 1953; Del Castillo and Katz, 1954b; Betz, 1970).

Understanding release requires a collective analysis of the events regulating vesicle fusion. An essential study by del Castillo and Katz (1956) showed that release may occur from any region of presynaptic terminals. It is also well-accepted that vesicle fusion requires a mature also called “primed” molecular assembly established by each vesicle with the plasma membrane (for review see Sudhof, 2013). Maturation of the fusion complex follows a stereotyped sequence of molecular transitions that can be resumed as follows: i. Docking (*D*) is the early tethering of vesicles with the plasma membrane upon establishment of boundaries between vesicle, membrane and soluble proteins; ii. Pre-priming (*pP*) occurs upon stabilization of the molecular complex; iii. Priming (*P*) occurs when vesicles become competent for fusion. Fusion (*F*) is evoked by calcium activation of the primed complex, mediated by the vesicle protein synaptotagmin. After fusion, vesicles are recycled to a new docked state (*F→D*) (del Castillo and Katz, 1956; Heuser and Reese, 1973; Betz and Angleson, 1998; Dulubova et al., 2005; Andrews-Zwilling et al., 2006; Kittel et al., 2006; Sudhof, 2013; Weimer et al., 2006; Imig et al., 2014; Gan and Watanabe, 2018). That only a small (~3%) fraction of the vesicle pool fuses on an impulse (Fatt and Katz, 1952, Katz and Miledi, 1979) has suggested that most vesicles rest in immature docked or pre-primed states.

Based on the stereotyped molecular transitions that render a primed vesicle and on the calcium-dependence of some such transitions (Neher and Sakaba, 2008; Burgoyne, 2007; Craxton, 2010; Corbalan-Garcia and Gómez-Fernández, 2014; Burgoyne et al., 2019), we put forward the hypothesis schematized in Figure 1, according to which the dynamic adaptations in the number of vesicles that fuse upon variations in nerve activity express a calcium-dependent, collective and reversible maturation of the fusion complex.

**Figure 1.**
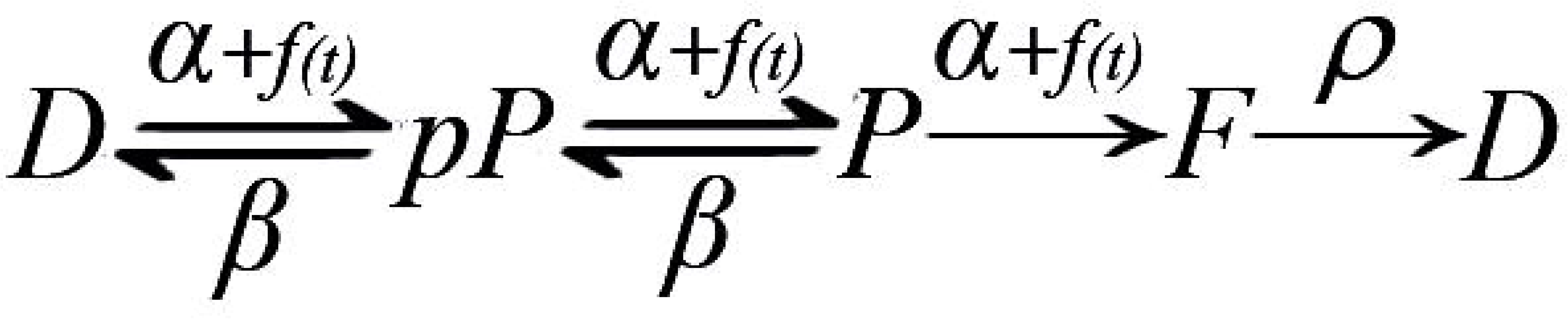
Kinetic model of molecular transitions of the fusion complex in individual vesicles. *D*, docked; *pP*, pre-primed; *P*, primed; *F*, fusion; *α*, forward rate constant; *f*_(*t*)_, calcium time dependence of the forward transition; *β*, backward rate constant; *ρ*, recycling rate constant. *D⇌ pP≠ P* are bidirectional; *P→F→D* are unidirectional; Spontaneous transitions occur following the corresponding rate constant; On electrical activity, the calcium-dependence accelerates the *Dp⇌P⇌P⇌F* transitions.

Our hypothesis considers that the *D*⇌*pP*⇌*P* transitions are bidirectional, with characteristic forward (*α*) and backward (*β*) rate constants. The *α* values are similar for all transitions; both *β* values are also similar but different from *α*. Reversibility is supported from the continuous docking and undocking of vesicles in ribbon synapses (Zenisek et al. 2000) and from experiments and modeling of pre-primed to primed transitions in crayfish neuromuscular junction (Pan and Zucker, 2009). On an action potential, calcium evokes fusion and promotes further maturation of fusion complexes. Rapid arrival of a subsequent impulse evokes facilitation. Backward transitions gradually reduce facilitation and return vesicles to resting levels. After intensive release, slow vesicle recycling new docked state plus the reversible transitions produce depression.

The experimental exploration of our hypothesis exceeds the current technical possibilities. However, mathematical modeling provides a solid alternative (Varela et al. 1997; Dittman et al. 2000; Shahrezaei, et al., 2006; Pan and Zucker, 2009; Dittrich et al., 2013; Herman and Rosemmund, 2015; Neher, 2015). Here we used a master equation based on the Gillespie (1976) stochastic algorithm to simulate the sequence of maturation transitions shown in Figure 1. Each vesicle with its fusion complex is a unit of a large homogeneous pool that responds collectively to each presynaptic impulse. The model was fed with experimental data from the literature. Undetermined parameters were fitted for convincing reproduction of well-known experiments of neuromuscular transmission in frog or cat. The code used for the simulations in this study is freely available in: https://github.com/alexini-mv/kinetic-neurotransmission

## Results

Spontaneous and evoked presynaptic vesicle fusion were accurately reproduced by a theoretical sequence of four maturation kinetic states of the vesicle fusion complex, provided that forward transitions had the same *α* value and were calciumdependent, while the backward transitions had a *β* value 50-100 times larger than *α*. A three-state model failed to reproduce transmission. By contrast, five or six sequential kinetic steps reproduced every form of release provided a proportional increase in *α* and reduction in *β*. The parameters that reproduced cat and frog neuromuscular transmission are resumed in Table 1.

**Table 1.** Kinetic parameters that reproduce neuromuscular transmission in frog and cat. * from Boyd and Martin, 1956a.

| Preparation | Kinetic transitions | $\alpha$ (s<sup>-1</sup>) | $\beta$ (s<sup>-1</sup>) | $\lambda$ ($\beta/\alpha$) | $\rho$ (s<sup>-1</sup>) |
| --- | --- | --- | --- | --- | --- |
| Cat | 4 | 0.62* | 62 | 100 | 1.0 |
| Frog | 4 | 0.3 | 15.0 | 50 | 1.0 |
| Frog | 5 | 0.62 | 13.0 | 21 | 1.0 |
| Frog | 6 | 1.43 | 9.5 | 13 | 1.0 |

### Spontaneous quantal release

The spontaneous quantal release in cat presynaptic neuromuscular junction reported by Boyd and Martin (1956a), was fairly reproduced by our model fed with an *α* = 0.62 s^−1^ value, obtained as the inverse of the experimental 1.61 s time constant (*τ*) of the time interval distribution of miniature end plate potentials (mepp_s_). An unexpectedly large *β* =100*α* (*λ*=*β/α* =100 coefficient) and a *ρ* =1.0 s^−1^ recycling rate constant produced 148 ± 2 meep_s_ at 1.40 ± 0.10 s^−1^ frequency (n=250 simulations), quite similar to the 143 meep_s_ recorded at a 1.43 ± 0.88 s^−1^ in the original paper (Figure 2A). The experimental distribution of the intervals between mepp_s_ was fitted by the function *n* = *n_T_*(Δ*t*/<*t*>) *e^−t/<t>^* (Fatt and Katz, 1952), where *n_T_* is the number of quanta released and Δ*t* = 0.5 s is the bin size.

**Figure 2.**
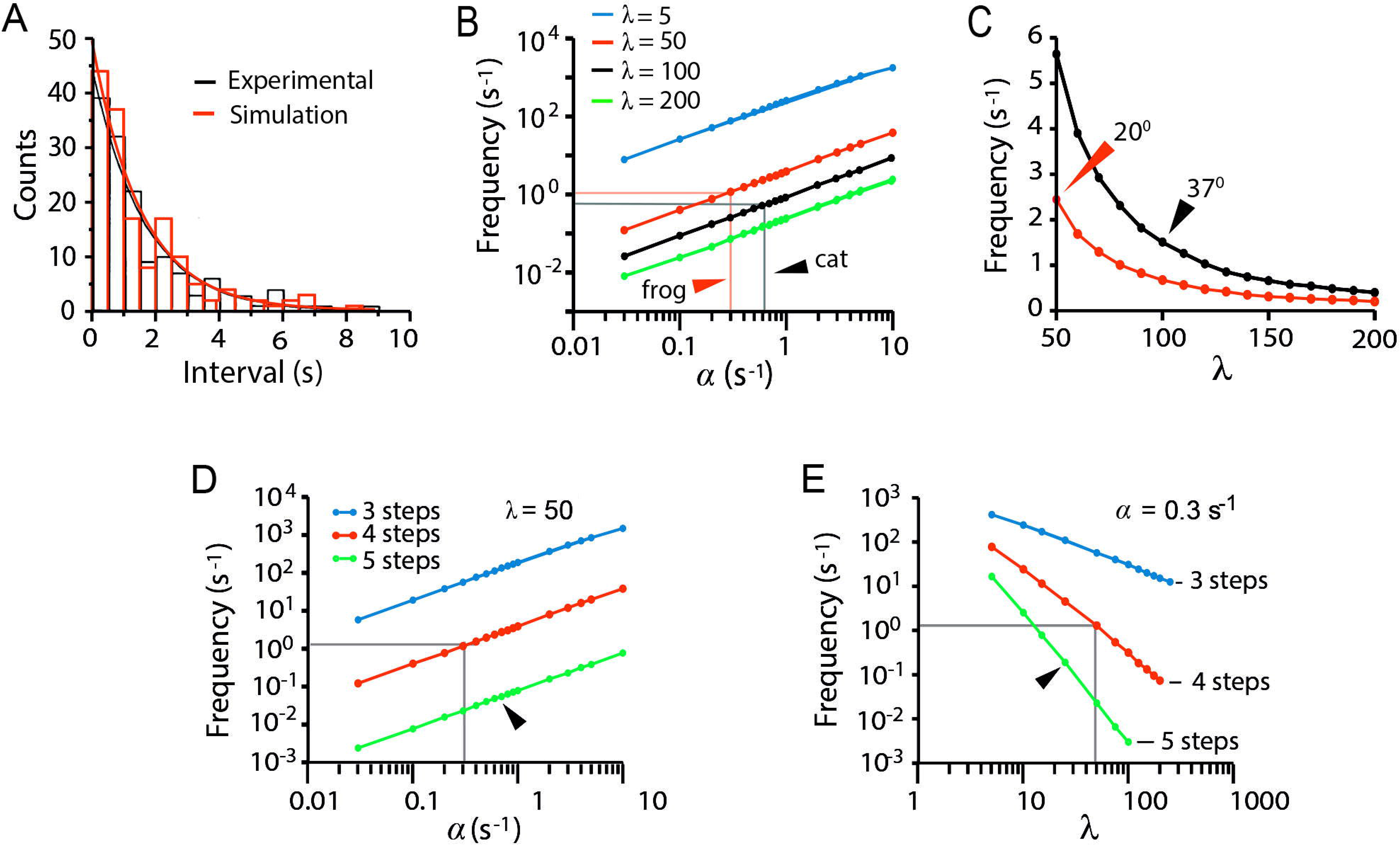
Spontaneous quantal release. A. Experimental (black) and simulated (red) time distributions of spontaneous mepp_s_ from 5-min recording intervals. The experimental distribution of 143 mepp_s_ was obtained with license from Boyd and Martin (1956b); the simulation contains 148 mepp_s_. The 1.54 s decay half time of the experimental probability rendered the *α*=0.62 s^−1^ value used in simulations of cat neuromuscular transmission along the paper. B) Predicted contributions of the *α* and *λ* values on the meep frequency. Arrowheads point to values that gave the best fittings in simulations of frog and cat transmission. C. Predicted mepp_s_ frequency as a function of the λ coefficient in frog (vermillon) and cat (black) synapses. The *ρ* = 1.0 value was equally successful in all simulations. Arrowheads point to experimental meep frequencies at the indicated temperatures, from Boyd and Martin (1956a). D. Effect of *α* on release with different number of kinetic steps. E. Effect of λ on release with different number of kinetic steps. The grey lines in D and E are the *α* and λ values that reproduce all forms of release in frog neuromuscular junction. Arrowheads indicate the *α* and λ values that reproduce release with 5 kinetic steps. Three kinetic steps failed to reproduce spontaneous release regardless on the *α* and λ values.

The meep_s_ frequency (Figure 2B) was proportional to *α* and inversely proportional to *β*. Our best explanation to this result is that the large *β* value reduces the pool of primed vesicles, therefore, the probability of spontaneous fusion. Figure 2C compares simulations of cat and frog spontaneous release. The *β* = 62 s^−1^ (*λ*=100) that reproduced the 1.43 s^−1^ meep frequency in cat recordings at 37°C (Boyd and Martin 1956a) quadruples the *β* =15 s^−1^ (*λ*= 50) coefficient that reproduced the 2.5 s^−1^ meep frequency commonly recorded from frog synapses at 20°C (see Fatt and Katz 1952). The larger rate constant values in mammalian neuromuscular synapses may reflect their characteristic higher physiological temperature.

The previous results may be explained in the following way. First, spontaneous release reflects the spontaneous *P→F* transitions; second, the small probability of spontaneous release depends on the large *λ* coefficient, which maintains a small pool of primed vesicles. Since a majority of experimental evidence used here proceeds from experiments in frog, the simulations that follow used the *α*=0.3 and *β*=50 values, unless otherwise indicated.

### Kinetic steps contributing to spontaneous release

Figure 2D and E shows that a three-step version of the model failed to reproduce spontaneous release. In addition, data in Figure 2D predicts that each kinetic step reduces the α-dependence of spontaneous release by two logarithmic units. By contrast, a five-step version of the model reproduced spontaneous release provided an increase of *α* (Figure 2D) and a reduction of *β* (Figure 2E). This means that a four-step *D⇌pP⇌P→F→D* transition cycle is necessary and sufficient to explain spontaneous release.

### Number of necessary kinetic steps for spontaneous release

Figure 2D and E shows that a three-step version of the model failed to reproduce spontaneous release. In addition, data in Figure 2D predicts that each kinetic step reduces the *α*-dependence of spontaneous release by two logarithmic units. By contrast, a five-step version of the model reproduced spontaneous release provided an increase of *α* (Figure 2D) and a reduction of *β* (Figure 2E). This means that a four-step *D⇌pP⇌P→F→D* transition cycle is necessary and sufficient to explain spontaneous release.

### Calcium and evoked release

A useful experimental strategy to study statistical fluctuations of quantal release consists of reducing the extracellular calcium concentration and adding extracellular magnesium (Del Castillo and Katz, 1954a; Boyd and Martin, 1955). Under such conditions, the number of quanta released by a presynaptic impulse is drastically reduced and can be predicted precisely by the Poisson distribution (Del Castillo and Katz, 1954a; Boyd and Martin, 1955). The theory states that the probability “*p*” of releasing “*x*” number of quanta (*x*=0, 1, 2, 3,…,*n*) in a series of trials is low, while the number “*n*” of vesicles in the pool remains large. Even when *p* and *n* are experimentally elusive, the product *m=pn*, which is the average number of quanta released per impulse is measurable from the recordings and provides a direct means for the calculations.

To reproduce such experimental observations, impulses were coupled to a “calcium elevation” whose amplitude and duration was adjusted to evoke release of small numbers of quanta (see methods). The hypothesis that nerve impulses induce forward transitions in each maturation was tested by coupling the calcium transient to every *α* rate constant. Based the observation by Katz and Miledi (1968 and 1979) that the amount of release increases with the duration of depolarization, i.e., with the duration and amount of calcium entry, we adjusted the decay time (*τ_e_*) of the artificial calcium signal as a way to control the amount of release. With such approximation the *m* value increased in proportion to *τ_e_*. The simulations in Figure 3A reproduce the experimental calcium-dependence according to the equation by Dodge and Rahamimoff (1966; see also Smith et al., 1985; Augustine and Charlton, 1986) expressing third (R^2^=0.999) or fourth order (R^2^=0.998) cooperativities.

**Figure 3.**
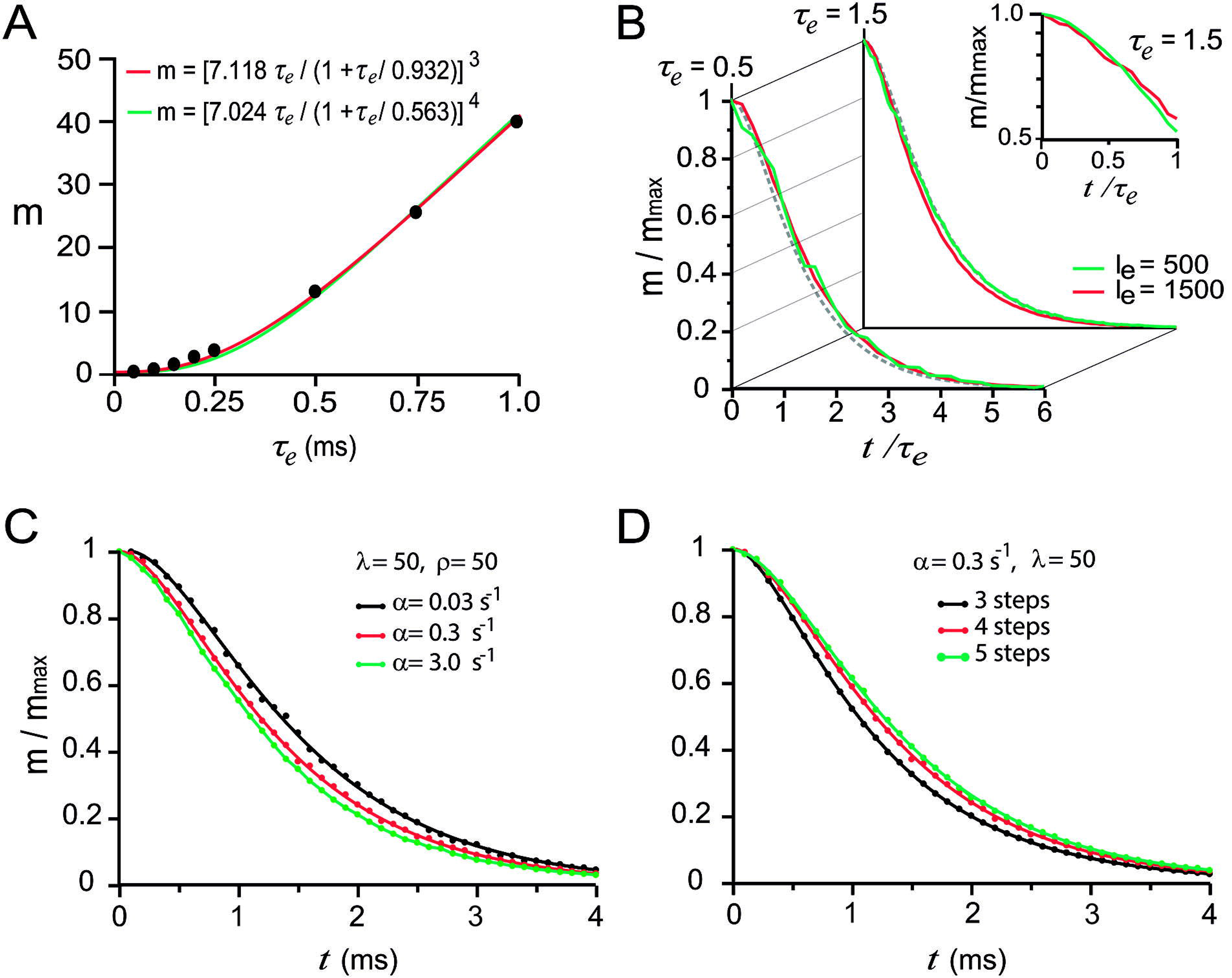
“Calcium-dependence” of quantal release. A. The mean number of quanta (m) depends on the mean decay time (τ_e_) of the intracellular calcium increase. The dots are model predictions; the lines were obtained with the equation by Dodge and Rahamimoff (1966) with third and fourth order cooperativities. B. The normalized number of quanta (m/m_max_) depends on the normalized *t/τ_e_* duration of the calcium signal. The traces are superimpositions of curves obtained using two different amplitudes (I_e_, arbitrary units) of calcium signal. The semilogarithmic chart in the inset shows the dispersion from a single exponential behavior below *t/τ_e_*=1. C. Increasing the *α* value accelerated release. D. Adding kinetic steps to the model increased the latency of release.

Another series of trials simulated calcium-dependent release under high release probability by using either, a long *τ_e_* value or a large calcium transient amplitude. The normalized number of quanta as a function of the normalized time (*t/τ_e_*) in Figure 3B displays a similar trend for different *τ_e_* and amplitude values. The combined effects of the amplitude and duration of the calcium transient in Figure 3B can be represented empirically by a sum of two exponential decays based on the hypothesis that kinetic transitions are represented by exponential equations in the form *m/m_max_* = (1 + *A*)*e^−t/τ_e_^* – *Ae^−t/xτ_e_^*. The detachment from a single exponential distribution originates from the second exponential component of the Equation, in which *x* expresses the combined contribution of *α* (Figure 3C) and the number of kinetic steps in the model (Figure 3D). Figure 3D shows that each step in the kinetic chain delays release presumably due to recruitment of vesicle fusion from every kinetic state. The lack of effect of *β* and *ρ* is attributed to the recovery of the vesicle pool between subsequent stimulation pulses.

### Evoked quantal release under low probability

Our model reproduced quantal release under low release probability in frog neuromuscular junction (Del Castillo and Katz, 1954a). Brief 0.05 to 0.15 ms *τ_e_* values produced mepp_s_ amplitude distributions with two (*τ_e_* = 0.05 ms) to five (*τ_e_* = 0.15 ms) amplitude classes including failures (Figure 4A). The Poisson equation reproduced such distributions when *τ_e_* < 0.5 (Pearson x^2^ > 0.05 coefficients). Larger *τ_e_* values produced a reduction in the number of failures and an increase in the number of classes in the distribution. *τ_e_* values longer than 0.25 produced amplitude distributions with marked deviations from the Poisson predictions (Figure 4B and C), as in experimental observations made under higher release probability (Boyd and Martin, 1956b).

**Figure 4.**
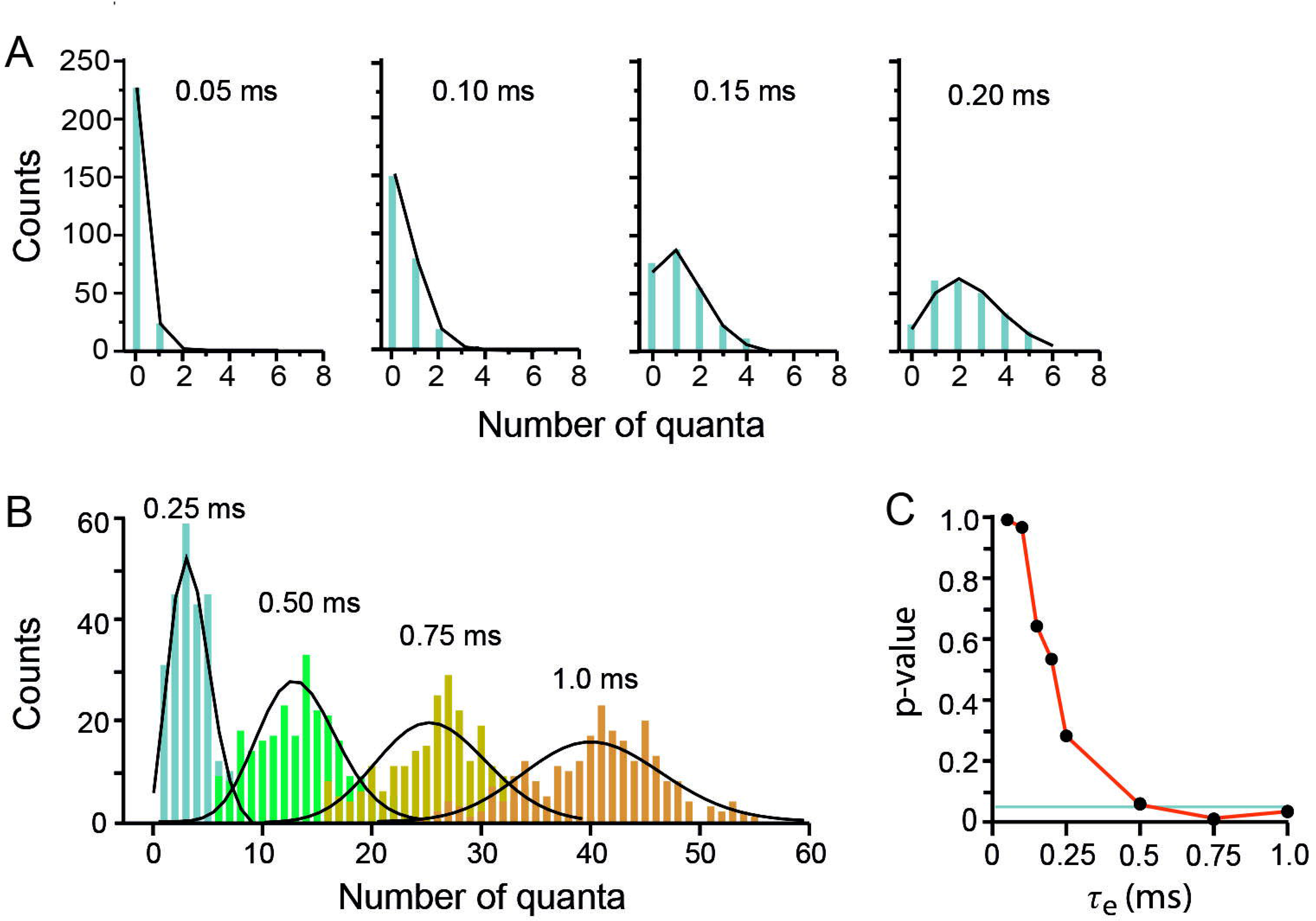
Evoked quantal release at low probability experimental conditions. A. Amplitude distributions of quantal release in frog neuromuscular junction. Counts are the number of quanta; the black lines link the discrete Poisson classes. The *τ_e_* values are above in each plot. B.-Amplitude distributions at increasing probabilities by use of larger *τ_e_* values. The discrepancies between the simulations and the Poisson predictions are clear with *τ_e_* values above 0.25 ms. Each plot contains data from 250 stimuli mediated by a 5 s recovery interval. C. Pearson’s significance (p) dependence on the *τ_e_* value. The horizontal line indicates the 0.05 significance.

### Role of the backward rate constant on the release probability

Simulations of frog experiments made under low probability conditions (Del Castillo and Katz, 1954a; Katz and Miledi, 1968) allowed a further analysis on the contribution of *β* to quantal release. The λ coefficients of the *D⇌Pp*(λ_1_) and *Pp⇌P*(λ_2_) transitions were varied independently, while the *α*=0.3 s^−1^, *ρ* = 1.0 and *τ_e_* = 0.15 ms remained fixed. The λ_1_=λ_2_ = 50 values reproduced transmission, as seen in the central chart of Figure 5.

**Figure 5.**
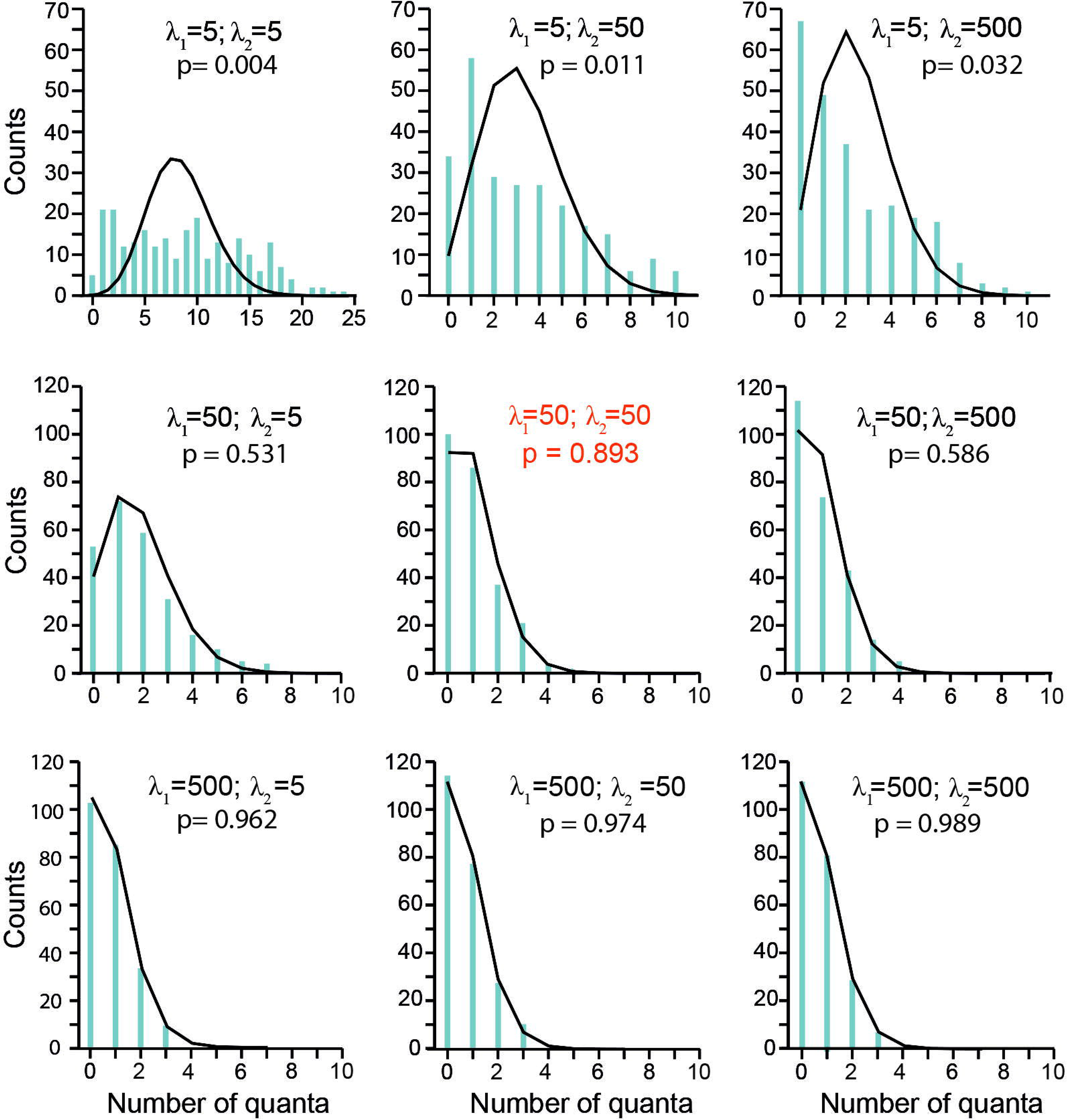
Contribution of the backward rate constant to quantal release. Data are presented in terms of the *λ=β/α* coefficients. *λ*_1_ corresponds to *D⇌pP*; *λ*_2_ corresponds to*pP⇌P*. The plots are as in Figure 4. The Pearson’s significance (p) appears in each chart. The central chart was obtained with λ_1_=λ_2_=50, which fitted every form of release in frog synapses. Other parameters were *α* = 0.3 s, *ρ* =1.0 s^−1^ and *τ_e_* = 0.15 ms.

The *λ*_1_ value markedly influenced the number of quanta discharged per impulse. A small *λ*_1_=5 (*β* = 5*α*; top panels in Figure 5) that decelerates the *D⇌Pp* transition extended the range of classes in the distribution, which deviated from the predictions of the Poisson equation (p≤ 0.05). Even the largest *λ*_2_= 500 tested failed to compensate for the affection caused a reduced *λ*_1_. By contrast, a large *λ*_1_=500 value constrained the amplitude meep distribution to a small-class range that was predicted by the Poisson distribution regardless of the *λ*_2_ value (bottom plots in Figure 5). As will be seen below, this result only applies to release under single impulses, for the large *λ*_1_=500 values failed to reproduce short-term plasticity. In spite of that, these results underscore the essential contribution of the backward *D⇌Pp* transition to maintain a small resting pool of primed vesicles.

### Facilitation and depression

Stimulation under experimental high release probability, for example by blocking acetylcholine receptors with curare (del Castillo and Katz, 1956; Betz, 1972), produces facilitation to turn into depression after a while, presumably because of depletion of the releasable vesicle pool (Otsuka et al., 1962; Mallart and Martin, 1968; Betz, 1970). In this section we reproduced the experimental transition from facilitation to depression in frog neuromuscular junction upon a conditioning train of three impulses followed by a test pulse applied 250 ms later, as in Mallart and Martin (1968). A *τ_e_* =1.3 ms reproduced the high release probability. The number of quanta, which is hard to estimate from experimental records, was predicted here by the model. An alternative approximation can be obtained from the relationship between the number of quanta and the *τ_e_* value in Figure 3.

Relevant to our study is that the same parameters that reproduce spontaneous and evoked release also reproduced the facilitation-depression balance. In addition, our simulations reproduced the asynchronous release that follows the bulk of evoked release on electric impulses in frog (Figure 6; Miledi, 1966) and other peripheral and central synapses (Zengel et al., 1980; Goda and Stevens, 1994; Atluri and Regehr, 1998; Hui et al., 2005; Best and Regehr, 2009).

**Figure 6.**
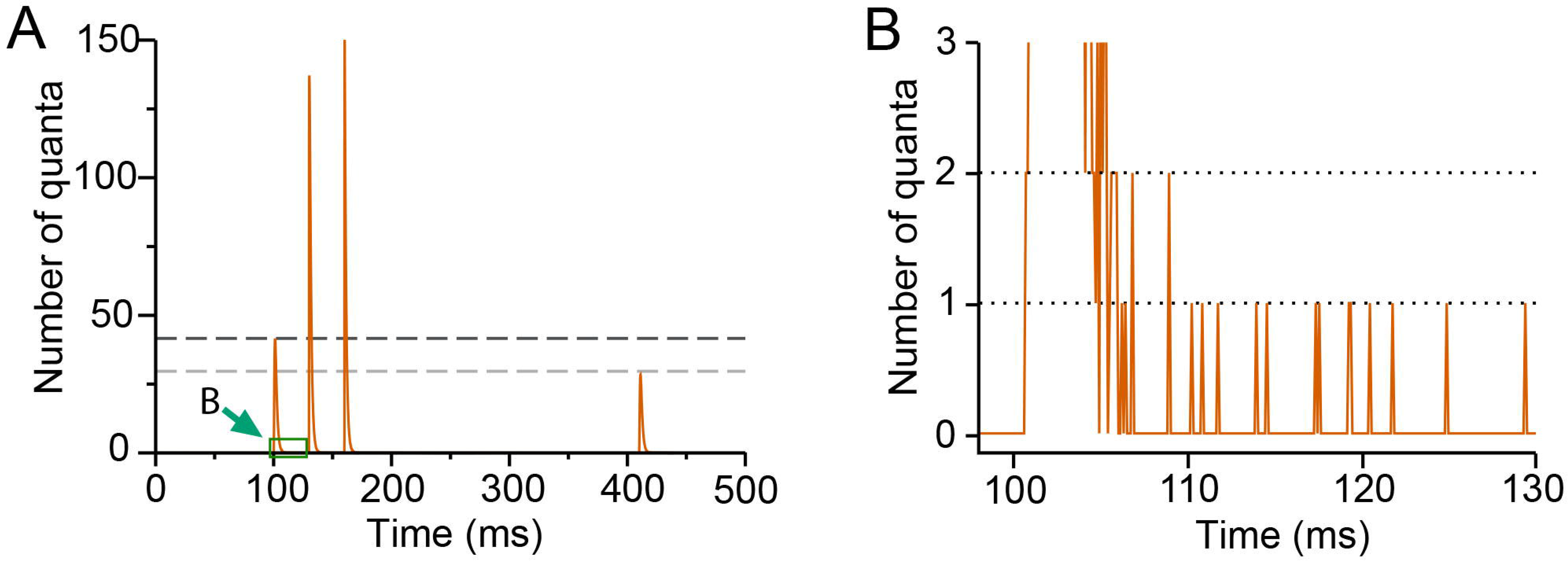
Sequence of facilitation and depression in frog neuromuscular junction. A) Number of quanta released in response to a train of three conditioning pulses 30 ms apart, followed by a test pulse 250 ms later (Mallart and Martin, 1968). Facilitation on the second and third impulses was followed by depression on the test pulse. B) Asynchronous release followed the conditioning impulses (square in A). The simulation parameters were *α* = 0.3 s^−1^; λ= 50; *ρ* = 1.0 s^−1^, and *τ_e_* =1.3 ms.

### The balance from facilitation to depression

How the sequence of transitions affects the balance from facilitation and depression in frog was analyzed with an alternative protocol used by Betz (1970). In this case, experiments were made with high extracellular calcium concentration to enhance depression and curare was added to block acetylcholine receptors and produce subthreshold transmission. Such conditions were reproduced here by *τ_e_*=1.5 ms (Figure 7). The rest were our conventional parameters for frog neuromuscular transmission. A briefer *τ_e_* =0.5 ms increased facilitation and abolished depression while a longer *τ_e_* =2.0 ms eliminated facilitation while depression remained.

**Figure 7.**
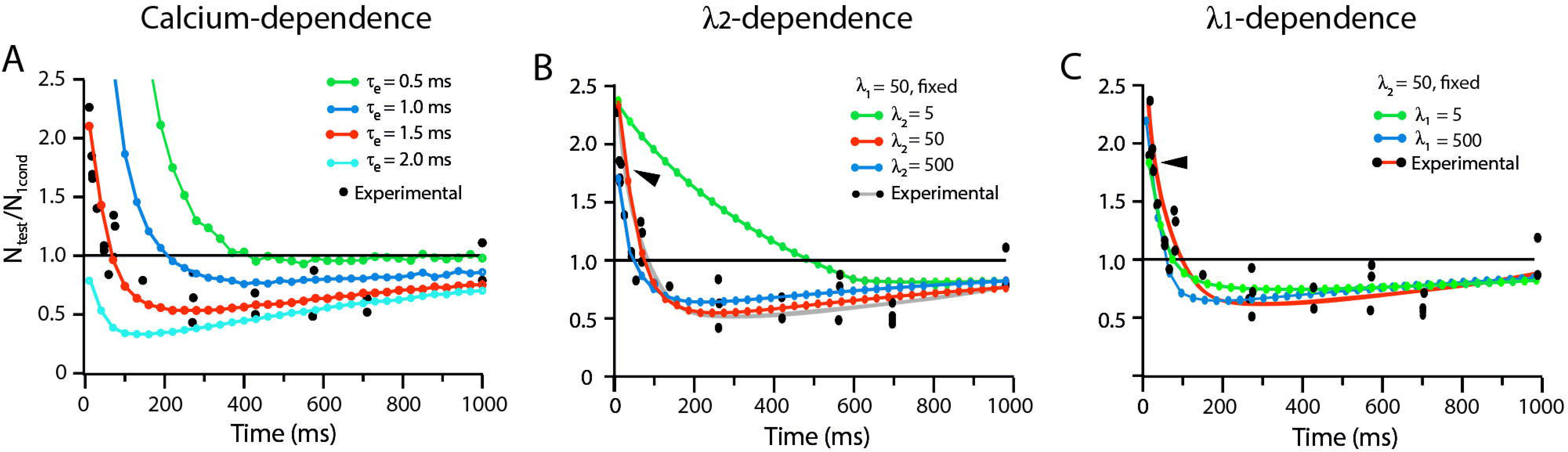
Calcium and rate constants influence short-term plasticity. A. The duration of the calcium signal (*τ_e_*)determines the balance between facilitation and depression. B. The λ_2_ coefficient determines the duration of facilitation. C. The *λ_1_* coefficient reduces facilitation and depression. N_test_/N_1cond_ is the ratio between the amplitude of the response to the test pulse (N_test_) and the conditioned pulse (N_1cond_). Values above 1.0 indicate facilitation; below 1.0 indicate depression. Experimental data obtained with license from Betz (1972).

### Effects of λ on short-term plasticity

Contrary to the dominant effect of *λ*_1_ on low probability release, the λ_2_ coefficient dominated facilitation (Figure 7B). A small λ_2_=5, which decelerates vesicle return to previous states elongated facilitation by 450%, from 90 to 500 ms, without affecting its peak amplitude. However, large λ_1_ = λ_2_=500 values reduced and shortened facilitation (arrowheads in Figure 7B and C). Increasing or decreasing any *λ* coefficient reduced depression without affecting its time course (Figure 7B and C).

### Effect of vesicle recycling on short-term plasticity

It has long been hypothesized that depression occurs when the releasable vesicle pool is reduced by large release and slow recycling dynamics (Elmqvist and Quastel, 1965; Kusano and Landau, 1975). The mild effects of λ on depression in our simulations support such hypothesis. Figure 8 shows that a ten-fold acceleration of the mean recycling time (*ρ* = 10 s^−1^) while keeping λ_1_ = λ_2_ = 50, increased the amplitude and duration of facilitation while depression was eliminated. Facilitation decayed biexponentially with a rapid *τ*_1_=30.19 ± 2.56 ms followed by a slower *τ*_2_ =169.55 ± 23.1 ms (R^2^=0.997). Conversely, a ten-fold reduction of *ρ* to slow down vesicle recycling had no effect on facilitation but increased depression from the N_test_/N_1cond_=0.6 in the experimental data to a 0.25 sustained value by 450 ms.

**Figure 8.**
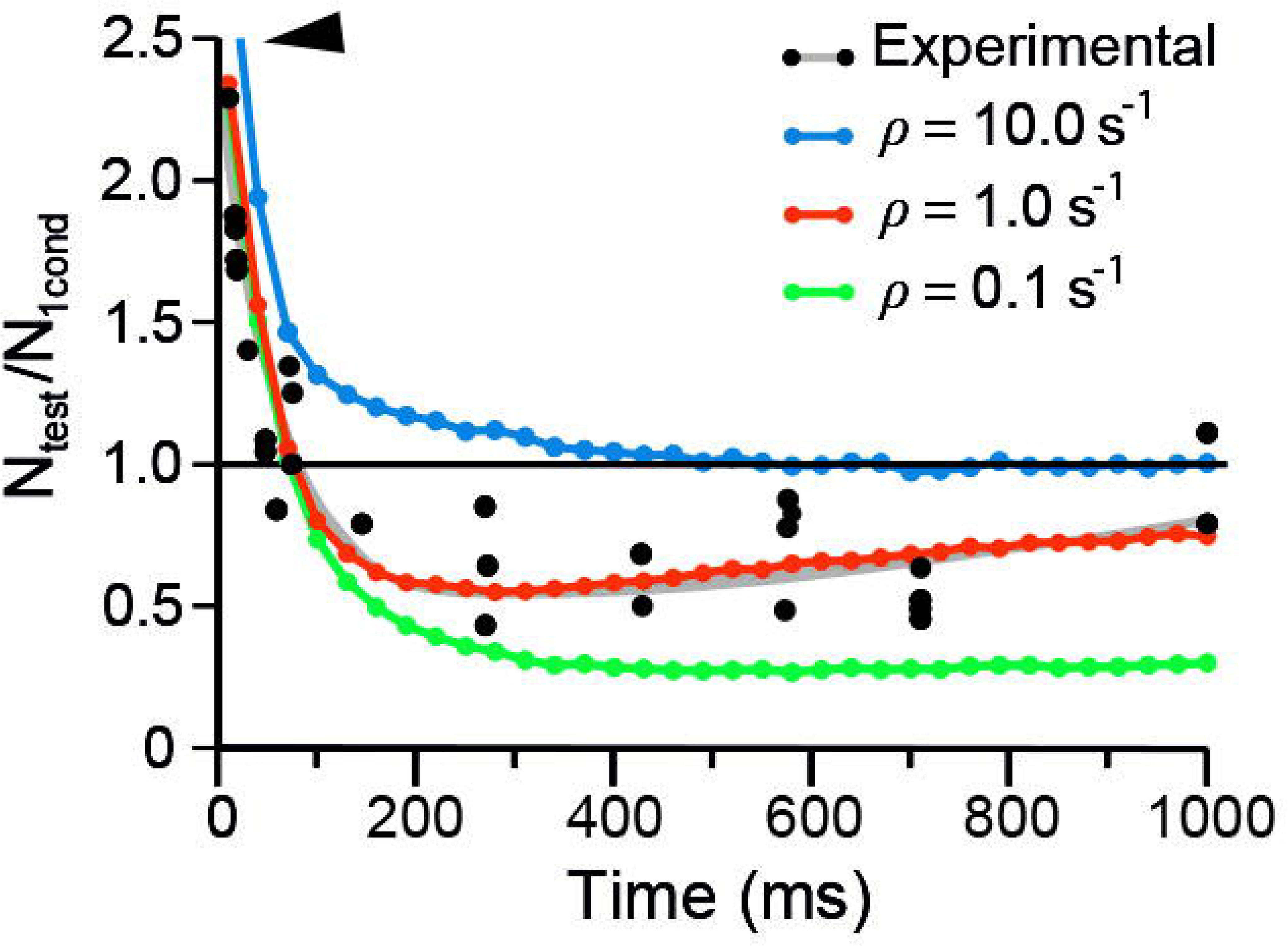
Vesicle recycling affects depression. The arrowhead denotes increased facilitation at high *ρ* value. Note the similar time course of the facilitation-depression sequence upon extreme *ρ* values. Conventional *α* and λ values for frog synapse were used.

### Effect of the number of kinetic steps on short-term plasticity

The three-step model fed with the regular parameters of frog experiments failed to reproduce facilitation but maintained depression levels similar to those already described (Figure 9A). By contrast, a five-step kinetic model by addition of a *D* state reproduced short-term plasticity provided a larger *α* = 0.62 s^−1^ (as in mammalian neuromuscular junction), and a reduced λ =21, accounting for a *β* = 13 s^−1^ value. Depression was less susceptible to the λ variations. A six-step model also reproduced the experimental data provided an even larger *α* =1.43 s^−1^ and smaller λ =13, reflecting *β* = 9.5 s^−1^. These results support that the four-state kinetic sequence fed with one common set of parameters is necessary and sufficient to reproduce the dynamics of release from spontaneous to short-term plasticity.

**Figure 9.**
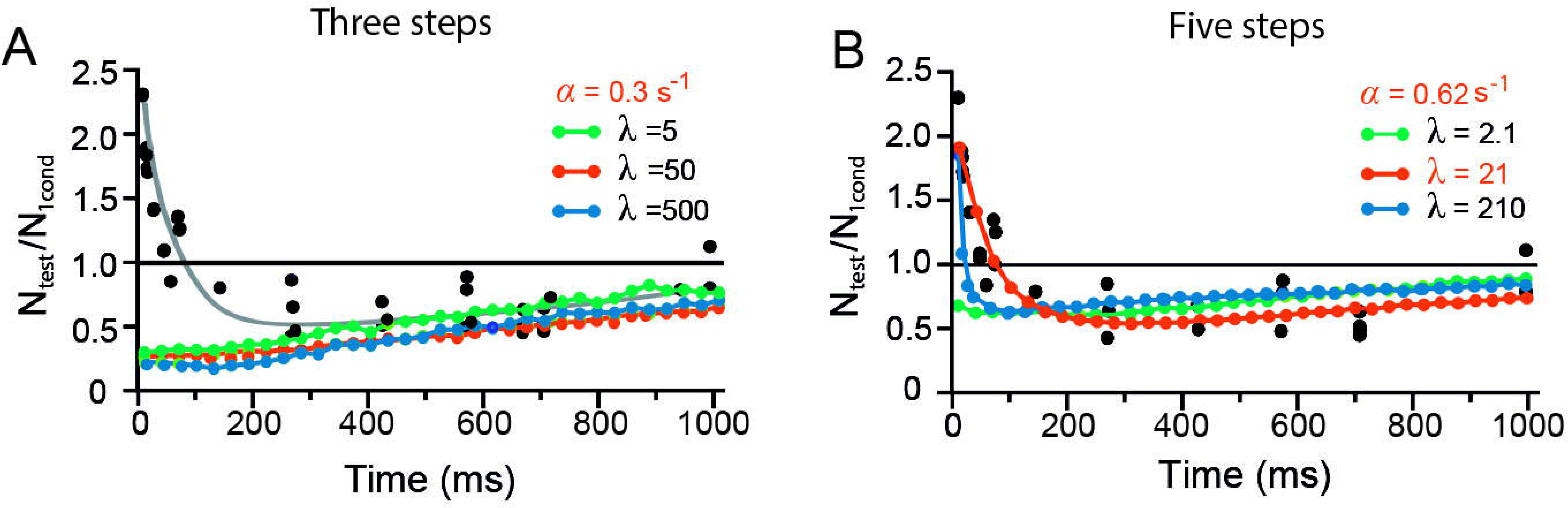
Short-term plasticity with different numbers of kinetic steps. A. Three kinetic steps produced depression without facilitation. B. Five kinetic steps reproduced short-term plasticity by using a larger *α* = 0.62 s and a smaller λ = 21.

### Activity-dependent dynamics of the vesicle pool

The plastic dynamics of transmission in response to a conditioning train followed by a test pulse were graphically plotted in Figure 10 (Figure 5; Mallart and Martin 1968). The fraction of vesicles in each state is normalized to N_0_=10,000. At rest, ~98% vesicles are docked, the remaining being decreasingly distributed in pre-primed and primed states. About 300 vesicles (3%) fuse in response to the first impulse, as estimated by Katz and Miledi (1979) at 6ū C. About 66% of the vesicles that fuse were primed, while the other 34% fuse upon rapid series transitions from immature states. The remaining vesicles become accommodated reversibly in immature states. Suden arrival of a second impulse encounters an increased population of pre-primed and primed vesicles, thus evoking facilitation plus further forward transitions. After the third conditioning pulse, about 25% of the vesicle pool has fused. Such large release along with the slow recycling (F/N_0_ panel in Figure 10) reduce the vesicle pool and produce a depressed response to the test impulse (Figure 10).

**Figure 10.**
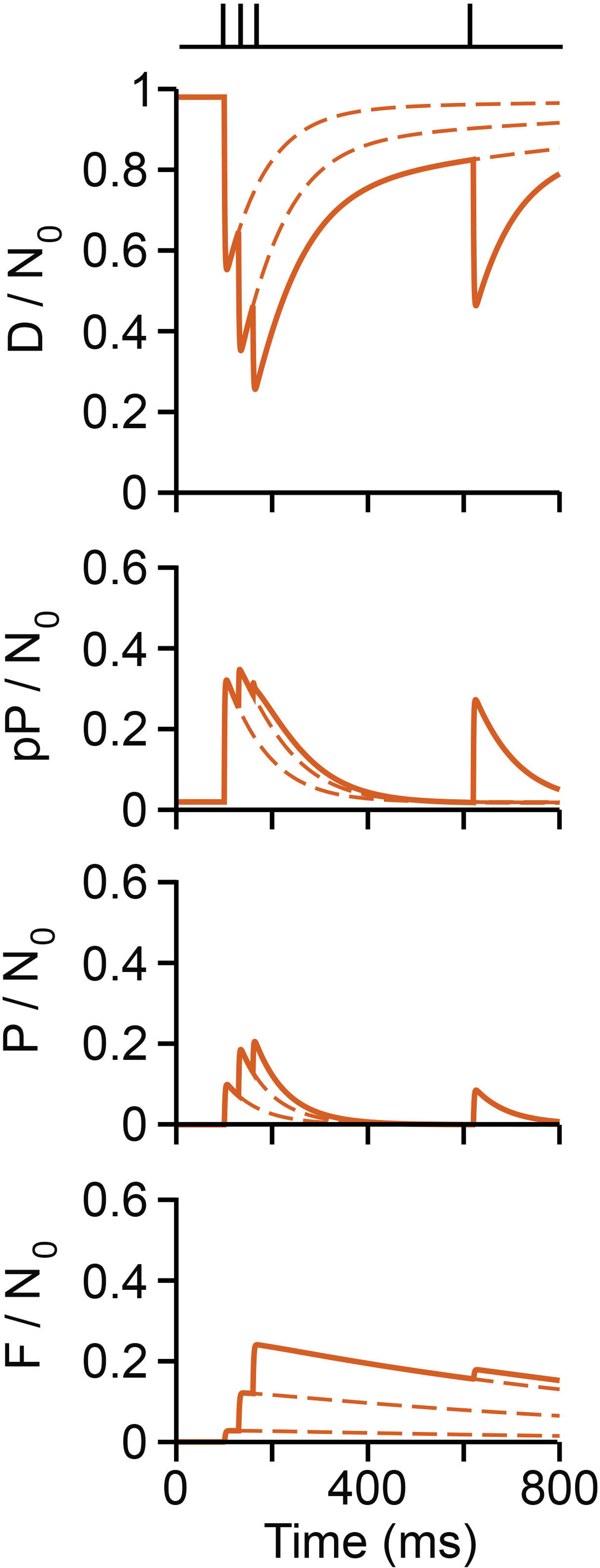
Vesicle dynamics in frog neuromuscular junction. The stimulation protocol is shown above (Mallart and Martin, 1968). The proportion of vesicles in each state is normalized to a pool of 10,000 (N_0_) vesicles. Calcium produces rapid *D⇌pP⇌P⇌F* transitions. Reversibility contributes to rapid recovery to resting state. Recycling contributes to slow recovery and depression.

## Discussion

Spontaneous release, evoked release and short-term plasticity, were reproduced here by a mathematical model of vesicles bound to a dynamic molecular fusion complex with four kinetic states. Our model provides a unifying mechanistic interpretation to the adjustable activity-dependent forms of release in a homogeneous vesicle pool. The backward rate constant and the much smaller forward rate constant values concentrate a vast majority of vesicles in docked state. Spontaneous and asynchronous fusion reflect spontaneous *P → F* occurrence in a small pool of primed vesicles. The dynamics of evoked release are influenced by electrical activity, which determines the momentary proportion of vesicles in the different maturation states. We also predict that the duration of facilitation depends largely on the backward transitions, with a contribution of the recycling time constant, whereas the duration of depression depends on the slow vesicle recycling.

### Functional advantage of the similar rate constant values

The accurate dynamics of transmission described here rely on collective transitions that result from the similar rate constants in each direction. Such similarity has been shown for the forward rate constants (Li et al., 2007; Chapman, 2008; Sudhof and Rothman, 2009). The synchronous collective transitions in either direction prevent the large accumulations of pre-primed and primed vesicles for long periods that occur when *λ*_1_ ≠ *λ*_2_. Such observation implies that the *pP* state is an essential buffer for adaptation of the vesicle pool of primed vesicles to the dynamics of motoneuron electrical activity. It was also shown that evoked release occurs from vesicles that were primed at the time of the impulse arrival with addition of vesicles reacting from immature states.

### Timing of facilitation and depression

The balance between the forward and backward transitions explains the timing of the non-linear fluctuations in the quantal output during facilitation and depression. The sequential transitions of the fusion complex produces that the pool of primed vesicles increases largely after the bulk of synchronous exocytosis. Therefore here, facilitation is due to the calcium-dependence delayed increase of pre-primed and primed vesicles on an impulse. Such delayed recruitment of previously immature vesicles is confirmed by the increased latency for fusion upon addition of a docked state to the model. A corollary to this observation is that the amount of facilitation relies on the balance between the duration of the calcium signal and the velocity of the forward transitions. Faster forward transitions would increase synchronous release and prevent facilitation. Slower forward transitions would produce small amounts of evoked release and rapid uncoupling of motoneuron and muscle responses.

### Relationship between facilitation and asynchronous release

Previous observations have suggested that facilitation and asynchronous neuromuscular release rely on the exact same mechanism (Rahamimoff and Yaari, 1973; Zucker, 1996). Our data suggest that such mechanism is the spontaneous fusion of primed vesicles. However, the dynamics of asynchronous release reflect the reversible activity-dependent maturation of vesicles from previous states.

### Recycling and depression

Our results confirm the essential role of vesicle recycling on depression and predict that backward transitions contribute to the amplitude of depression. Two or more recycling modes in the neuromuscular junction (Rizzoli and Betz, 2005) and central synapses (Wu and Borst, 1999; Sakaba and Neher, 2001) suggest equal numbers of recycling vesicle pools (for review see Alabi and Tsien 2012). However, with a single recycling rate constant, our model reproduced convincingly the balance between facilitation and depression as studied by Betz (1970). However, we cannot exclude that the slow time constant of recycling in our model is masking faster events including some displaying a calcium-dependence (Sakaba and Neher, 2001).

### A single type of calcium sensor for fusion

Our data are consistent with one type of calcium sensor producing all forms of release, in contrast to the proposed contribution of different forms of synaptotagmin controlling release in central synapses (Volynski and Krishnakumar, 2018; Sudhof, 2013; Keser and Regher, 2014; Kavali, 2015). Central synaptic vesicles seem to carry different types of synaptotagmin (Jahn and Sudhoff, 1994; Takamori et al., 2006), and fast synchronous release is produced by synaptotagimnes 1, 2 or 9 (For review see Kaeser and Regehr, 2014) asynchronous release depends on the high calcium affinity synaptotagmin 7 (Bacaj et al., 2013, 2015; Turecek and Regehr, 2018). Accordingly, theoretical models of transmission considering two or three calcium sensors reproduce the electrophysiological data (Dutta Roy et al., 2014; Goda and Stevens, 1998). Although neuromuscular terminals of the genetically-accessible Drosophila, zebrafish and mice express synaptotagmins 1 and 2 (Pang et al., 2006), careful experiments in Drosophila neuromuscular junction have shown that synaptotagmin 1 mutants cannot evoke fast release (Shields et al., 2020), supporting that synaptotagmin 2 is functionally enough for fast fusion, in consistency with our results.

### Calcium-sensors for early transitions

In addition to the catalyzing role of calcium on vesicle fusion, we predict that calcium also promotes the *D → pP → P* transitions. A natural question emerging from such hypothesis is if the same calcium sensor producing fusion may also produce the previous transitions. The residual calcium hypothesis for paired pulse facilitation by Katz and Miledi (1968) and the third or fourth order calcium-dependence of release (Dodge and Rahamimoff, 1966; Smith et al., 1985; Augustine and Charlton, 1986) predict that the low residual calcium levels activate high-affinity calcium sensors to produce the supralinear vesicle fusion in facilitation (Zucker and Lara Estrella, 1983; Yamada and Zucker; 1992; Van der Kloot and Molgó, 1993; Vyshedskiy and Lin, 1997; Regehr and Zucker, 2002; Ma et al., 2014). Such high affinity sensor must be other than the low affinity synaptotagmins 1, 2 or 9 (For review see Kaeser and Regehr, 2014). Synaptotagmin 7 has emerged again as a candidate in central synapses (Sugita et al., 2002; Pan and Zucker, 2009; Wen et al., 2010; Bacaj et al., 2013, 2015; Jackman and Regehr, 2014; Li et al., 2017; Turecek and Regehr, 2018). However, the possibility of a single sensor producing all serial transitions from pre-priming to fusion is feasible in neuromuscular synapses by considering that electron tomography experiments have shown that from the moment of docking, the fusion complex forms intimate boundaries with calcium channels (Harlow et al., 2001; Nagwaney et al, 2009; Szule et al., 2012). Such configuration may permit a single calcium sensor to catalyze every transition of the chain, as opposed to central synapses in which calcium channels may be separated from the fusion complex at early maturation stages (Neher, 2015).

## Methods

### Design of the mathematical model

The four-state kinetic model with six kinetic transitions shown in Figure 1 set the bases to analyze the collective behavior of a pool of 10,000 identical vesicles (Rizzoli and Betz, 2005). The six transitions *R_j_*, correspond to those in Figure 1, with *j* being a stochastic discrete variable that takes the values *j=1, 2,…6*, that correspond to each transition. Each transition occurs with an equal probability *a_j_*(*x*). The term *a_j_*(*x*)*dt* is the probability that an *R_j_* transition will occur in an infinitesimal time interval *t+dt*, when the system is in a state *X(t)=(D(t), pP(t), P(t), F(t))=x*. Each *R_j_* transition is characterized by two quantities: One is the system state *x* = *D*(*t*),*pP*(*t*),*P*(*t*),*F*(*t*), which reflects the number of vesicles at each kinetic state; the second quantity is the vector *V_j_*(*v_D_j__, v_pP_j__, v_P_j__, v_F_j__*), which represents the change in the total number of vesicles over time at each state. At rest, a vast majority of vesicles lay in the *D* state. The effect of larger numbers of molecular states on transmission was analyzed by adding states with their corresponding bidirectional rate constants, between the *D* and *pP* states. For a three-state possibility the*pP* state was eliminated from the model.

The stochastic kinetic model considers that fusion requires vesicles to arrive at the *P* state. Since the classical kinetic differential equations do not describe correctly the collective kinetics of a small number of vesicles (~10,000 as compared to Avogadro’s number), we used instead the master Equation (1) for the probability distribution *P*(*x,t;x_0_,t_0_*) (Gillespie, 1976), whose solution describes the temporal evolution of the six transitions between kinetic states. The rate constants are conventional probabilities per time unit (Gillespie, 1992):

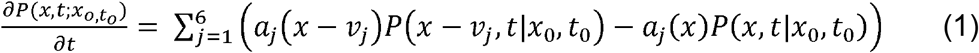

Because calculating an analytical solution of Equation 1 in extremely difficult, the solution of Equation 1 was obtained using the Gillespie algorithm (Gillespie, 1976), which emulates random transitions connecting different *X*(*t*) states. The fundamental equation of the Gillespie algorithm for the time evolution of the system is:

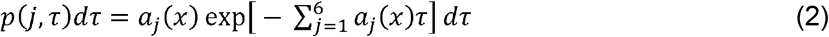

Equation 2 predicts the probability that at a state *X(t) = x*, the next kinetic transition *R_j_*, will occur at the next infinitesimal time [*t + τ,t + τ + dτ*]. The random continuous variable *τ* advances the time in the simulations by the amount:

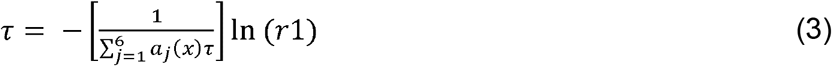

with *r*1 being a random number distributed uniformly in the interval (0, 1).

The probability distribution *p*(*j,τ*)*dτ* mimics the solution of the stochastic kinetic Equation 1 and plays a key role in the implementation of the stochastic algorithm. Thus, the random trajectories that connect different kinetic states, *X*(*t*) = *x*, describe the kinetic evolution of the vesicle pool.

The algorithm for the kinetic sequence can now be summarized as follows: (i) The simulation begins by setting the initial state of the system *X_o_* at time *t_o_*. (ii) The propension functions *a_j_*(*x*) and their sum *a_o_*(*x*) = *∑aj*(*x*) are calculated for each different time t. (iii) The values of the discrete random variables *j* is chosen as the smallest integer that satisfies, 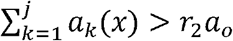 with *r*_2_ a random number distributed uniformly in the interval [0,1], and the continuous random variable *τ* is generated by applying equation (3). (iv) The transition to the next kinetic state *x → x + v_j_* and the time shifts to *t → t + τ* are calculated. (v) A new state (*x, t*) is obtained, and the procedure returns to step (i).

The simulation starts with *No* =10,000 vesicles accumulated in the *D* state. In such conditions *X(t = 0) = Xo = (D(t = 0) = No*, and *pP(t = 0) = 0, P(t = 0) = 0, F(t = 0) = 0)*. As the simulation progresses, the vesicles distribution among the different states becomes stationary in about 5 min. This is the time in which our measurements in the simulations are made.

### Estimates of kinetic values

The activation energies involved in the molecular transitions from docking to exocytosis lay in the same order of magnitude (Li et al., 2007; Sudhof and Rothman, 2009). Therefore, we initially considered that *α*_1_=*α*_2_=*α*_3_=*α*, and *β*_1_=*β*_2_=*β*. Such strategy proved successful for reproducing every release mode. The *α* value used in cat simulations was estimated from the frequency distribution of spontaneous miniature potentials (Boyd and Martin, 1956a, b). The *β* and *ρ* values were fitted independently. The model was simplified by using the coefficient *λ=β/α* which permitted to evaluate the kinetic behavior in terms of the combination of *ρ* and the relative magnitudes of *α* and *β* The code used in this study is available in the following repository: https://github.com/alexini-mv/kinetic-neurotransmission

### Modeling the calcium-dependence

Presynaptic calcium elevations upon brief depolarizations were modeled by adding a function *f*(*t*) to the forward rate constants, which acquired the form *a*_s_ = *α* + *f*(*t*). The kinetics of the calcium current decay in squid giant synapse experiments (Llinas et al., 1981a and b) served as baseline. The onset of the calcium transient was considered as instantaneous for the calcium channels in presynaptic neuromuscular terminals are tightly bound to the fusion complex (Harlow et al., 2001; Nagwaney et al, 2009). Adjustments in the amplitude (in arbitrary units) and decay time (ms) of the artificial calcium elevation rendered successful results.

For our simulations it was more convenient to express the decay time *τ_e_* of the calcium elevation instead of the decay time of the current, since according to the residual calcium hypothesis (Katz and Miledi, 1968; Kamiya and Zucker, 1994; Matveev et al., 2006), it is the remnant free intracellular calcium after the impulse what promotes facilitation. The decay time of the calcium elevation was defined as:

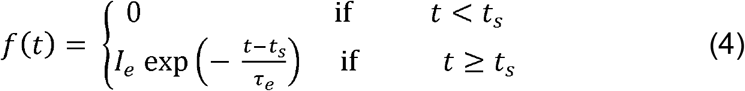

where *t_s_* is the stimulation time. The *τ_e_* value was adjusted for each experimental protocol in a range of 0.05 - 1 ms. Once adjusted, the parameters of the calcium signal remained the same for each experiment. Calcium currents in certain central synapses may facilitate or depress upon subsequent stimulation (Borst and Sakmann, 1998; Forsythe et al., 1998; Cuttle et al., 1998; Inchauspe et al., 2004; Ishikawa et al., 2005; Xu and Wu, 2005; Mochida et al., 2008). However, our model rendered accurate results without any such modulation.

## Acknowledgements

We wish to express our gratitude to Mr. Bruno Mendez and to Mrs. Sara Flores González for their excellent laboratory assistance. Our research was funded by a DGAPA-UNAM grant IN200914 and a CONACYT grant 130031 to FFDM and by a DAGAPA-UNAM grants IN118410 and IN108916 to GRS. AMV acknowledges support from CONACYT as a master’s degree fellowship at Posgrado en Ciencias Físicas at UNAM.

